# Biosynthesis- and Metabolomics-guided discovery of antimicrobial cyclopeptides against drug-resistant clinical isolates

**DOI:** 10.1101/2023.10.26.563470

**Authors:** Zhuo Cheng, Bei-Bei He, Kangfan Lei, Ying Gao, Yuqi Shi, Zheng Zhong, Hongyan Liu, Runze Liu, Haili Zhang, Song Wu, Wenxuan Zhang, Xiaoyu Tang, Yong-Xin Li

**Affiliations:** Department of Chemistry and The Swire Institute of Marine Science, The University of Hong Kong, Pokfulam Road, Hong Kong, China; Institute of Molecular Chemical Biology, Shenzhen Bay Laboratory, Shenzhen, China; State Key Laboratory of Bioactive Substance and Function of Natural Medicines, Institute of Materia Medica, Chinese Academy of Medical Sciences and Peking Union Medical College, Beijing 100050, China

## Abstract

Antimicrobial resistance remains a significant global threat, contributing significantly to mortality rates worldwide. Ribosomally synthesized and post-translationally modified peptides (RiPPs) have emerged as a promising source of novel peptide antibiotics due to their diverse chemical structures. Here, we reported the discovery of new Avi(Me)Cys-containing cyclopeptide antibiotics through a synergistic approach that combines rule-based genome mining, automated metabolomic analysis, and heterologous expression. We first bioinformatically identified 1,172 RiPP biosynthetic gene clusters (BGCs) responsible for Avi(Me)Cys-containing cyclopeptides from a vast pool of over 50,000 bacterial genomes. Subsequently, we successfully established the connection between three newly identified BGCs and the synthesis of five new peptide antibiotics. Notably, massatide A displayed excellent activity against a spectrum of gram-positive pathogens, including drug-resistant clinical isolates like linezolid-resistant *S. aureus* and methicillin-resistant *S. aureus*, with a minimum inhibitory concentration (MIC) of 0.25 μg/mL. The remarkable performance of massatide A in an animal infection model, coupled with a low risk of resistance and favorable safety profile, positions it as a promising candidate for antibiotic development. Our study highlights the potential of Avi(Me)Cys-containing cyclopeptides in expanding the arsenal of antibiotics against multi-drug-resistant bacteria, offering promising drug leads in the ongoing battle against infectious diseases.

## Introduction

The rise of multi-drug-resistant bacteria, commonly called “superbugs,” has become an alarming global health concern, leading to untreatable infections^1^. The urgent need for new antibiotics to combat these superbugs is evident. In addition to conventional small-molecule antibiotics, scientists have turned their attention to antimicrobial peptides as a potential solution. Among them, macrocyclic peptides, including nonribosomal peptides (NRPs) and ribosomally produced post-translationally modified peptides (RiPPs), have emerged as a promising source for antibiotic discovery^2^. Their favorable properties, achieved through macrocyclization, enable them to inhibit proteolysis and maintain an active conformation for effective target binding^3^. Novel RiPP antibiotics, such as ruminococcin^4^, thiostrepton^5^, nosiheptide^6^, darobactin^7^ and dynobactin^8^, as well as NRPs, including teixobactin^9^, brevicidine^10^, rimomycin^11^, misaugamycin^11^, corbomycin^12^, and clovibactin^13^, highlights the potential of macrocyclic peptides in fighting bacterial infections.

However, traditional methods of discovering new antibiotics have proven inadequate, leading to a shortage of novel antibiotics to combat increasingly resistant bacteria worldwide^1, 2^. Advances in sequencing techniques have enabled large-scale sequencing of microbial genomes, opening new opportunities for discovering novel peptide antibiotics through genome mining^14^. Among the vast array of natural products, RiPPs stand out as one of the most expansive families, offering a remarkable diversity of structures and bioactive potential for antibiotic discovery^15^. Consequently, significant efforts have been devoted to genome mining-guided discovery of new RiPP antibiotics^16, 17^, taking advantage of the genetically encoded nature of RiPPs to explore their chemical space and predict their chemical structures efficiently. In recent studies, our research focused on genomics-guided discovery of RiPP antibiotics, resulting in the discovery of antagonistic lanthipeptide from archaea and narrow-spectrum class II bacteriocins from the human microbiome^18, 19^. Moreover, applying rule-based genome mining strategies, we successfully revealed previously hidden peptidases^20^ and untapped post-translational modification (PTM) enzymes^21, 22^ for RiPP antibiotic biosynthesis. However, the challenge remains in the prioritization of BGCs of interest and the robust linking of metabolites to their corresponding BGCs prior to time-consuming isolation and identification. Here, our continued endeavors are to access untapped RiPP biosynthetic potential through combinational usage of rule-based genome mining, automated metabolomic analysis, and heterologous expression, effectively linking them to chemicals to discover peptide antibiotics.

Aminovinyl-(methyl-)cysteine-containing cyclopeptides (ACPs), which possess structurally distinctive cyclic peptide rings and antibacterial activity, have recently emerged as promising candidates in these endeavors^23^. The rigidity of the Avi(Me)Cys unit restricts the conformational flexibility of the peptides, inhibiting proteolysis and maintaining the peptide in an active conformation for target binding^23^. This property confers multiple bioactivities to ACPs, as demonstrated by microbisporicins, which interact electrostatically with the negatively charged lipid II pyrophosphate bridge, making them effective against vancomycin-resistant bacteria^24, 25^, and lexapeptide, which exhibits potent bioactivity against methicillin-resistant *Staphylococcus aureus* (MRSA) and methicillin-resistant *Staphylococcus epidermidis* (MRSE)^26^. To date, the Avi(Me)Cys moiety has been found in five RiPP families, including lanthipeptides, lipolanthins, lanthidins, thioamitides, and linaridins^23, 27–32^ (Figure 1a, S1). The genetically encoded nature and diverse biosynthesis pathways of ACPs allow for the efficient expansion of their chemical space. Together with their unique structural features, promising bioactivities, and the increasing availability of microbial genomes, we are motivated to conduct a systematic bioinformatic investigation of underexploited ACPs for the discovery of new antibiotics.

**Figure 1.**
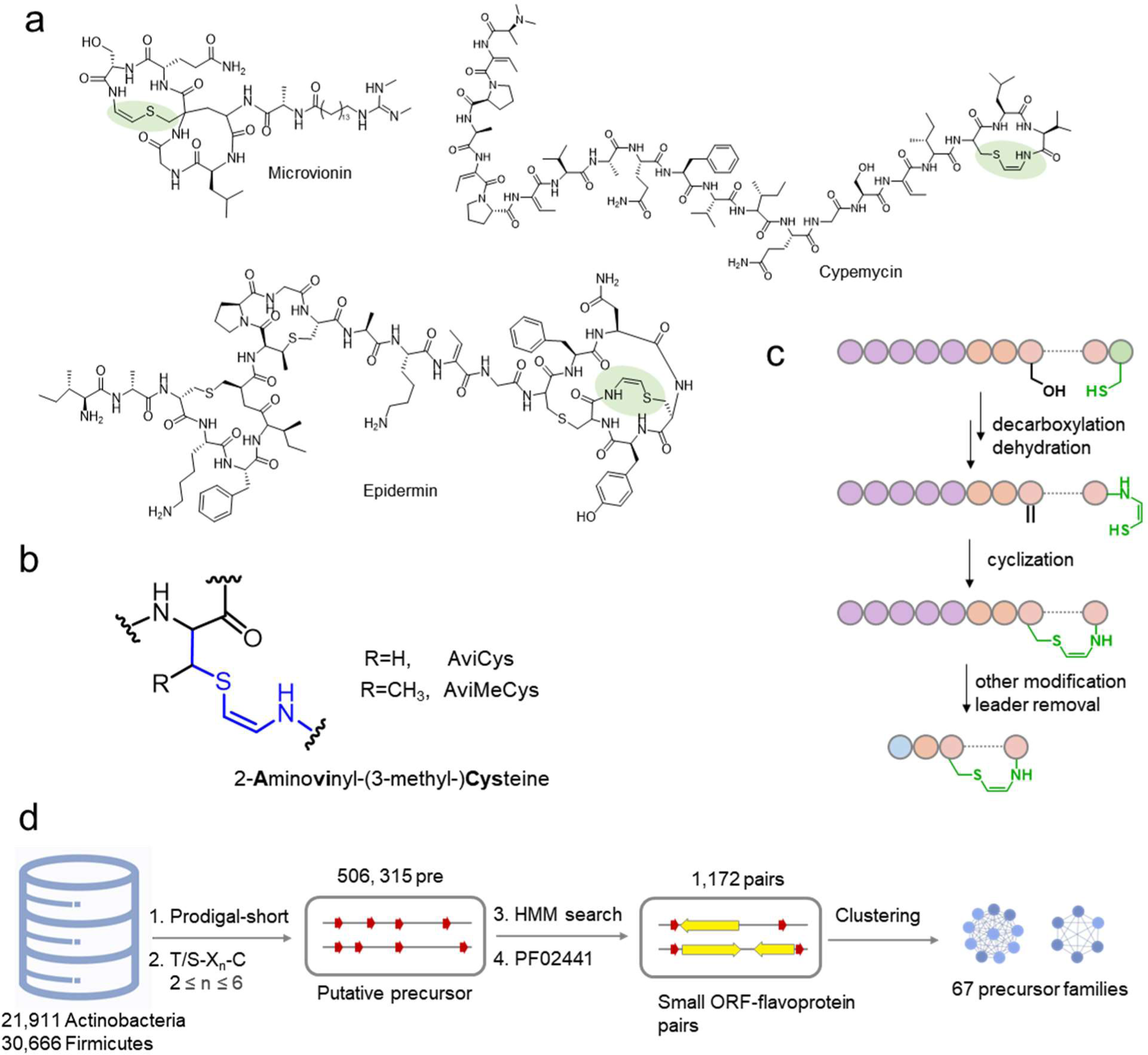
Representatives, biosynthesis and genome mining of aminovinyl-cysteine-containing peptides. a) Representative examples of Avi(Me)Cys-containing RiPPs, with the Avi(Me)Cys motifs highlighted in green. b) The structure of Avi(Me)Cys motif. c) A common biosynthetic process for Avi(Me)Cys unit formation. d) Genome mining pipeline of Avi(Me)Cys-containing RiPP BGCs using SPECO.

Despite the structural diversity of ACPs resulting from various modifications, such as the methylation in linaridin^27^, fatty acid chain in lipolanthin^30, 31^ and thioamide bond in thioamitide^28, 33^, the biosynthetic machinery towards the Avi(Me)Cys unit is highly conserved (Figures 1b,c, S1). A characteristic flavoprotein catalyzes the oxidative decarboxylation of precursor C-terminal cysteine to yield enethiol, followed by thioether cyclization with dehydroalanine (Dha) or dehydrobutyrine (Dhb) to generate AviCys or AviMeCys^23^ (Figure 1c). Building on this conserved biosynthetic logic, we conducted a large-scale analysis of publicly available actinobacteria and firmicutes genomes using the rule-based genome mining pipeline SPECO^21^ (Figure 1d). Through sequence similarity network (SSN) and sequence logo analysis, we discovered 1,172 BGCs, including 67 ACP families. By implementing an automated mass mapping pipeline, we link five ACPs to their corresponding BGCs prior to isolation, further supported by the successful heterologous expression of BGCs in a streptomyces host. Furthermore, our bioactivity assay demonstrated that the newly discovered ACP antibiotics exhibit remarkable potency against gram-positive bacteria, as evidenced by their strong inhibitory activity both *in vitro* and *in vivo*. These discoveries open up exciting prospects for identifying and characterizing novel ACPs, which could serve as a valuable source of new antibiotics to combat the increasing prevalence of multi-drug-resistant bacteria.

## Results

### SPECO-based genome mining expands Avi(Me)Cys-containing RiPP families

RiPPs are biosynthesized using a conserved mechanism whereby post-translational modification (PTM) enzymes modify a ribosomal precursor peptide, followed by protease-mediated trimming to release the mature product^15^. Similarly, ACP biosynthesis relies on a flavoprotein (Pfam number: PF02441) that catalyzes the oxidative decarboxylation of the C-terminal cysteine residue of the precursor. This process results in the formation of an enethiol group, which can react with unsaturated amino acids, facilitated by oxidative thioaldehyde formation of the cysteine residue and parallel decarboxylation through tautomerization^23^. The requirement of the flavoprotein for thioenol formation and the indispensability of the C-terminal cysteine residue in precursor peptides provide a unified rule for genome mining of ACP-encoding BGCs (Figure 1c).

Following this rule, we analyzed 21,911 genomes of actinobacteria and 30,666 genomes of firmicutes for the ACP BGC identification using the SPECO pipeline^21^ (Figure 1c). This analysis generated 1,172 unique putative precursor-flavoprotein pairs (Supplementary dataset). Our findings indicate that most of the putative precursors were not clustered with known precursors (Figure 2a) in the sequence similarity network (SSN), revealing the unexplored chemical space of ACPs. As expected, known precursors such as epidermin, thioviridamide, lexapeptide, and cypemycin precursors were included in the top-rank (Figure 2a). We further analyzed the putative BGCs through sequence logos and phylogenetic analysis to prioritize candidates with unknown precursors or tailoring enzymes (Supplementary dataset). Among the identified BGC families, one family caught our attention due to the presence of two oxidoreductases, which is uncommon in ACP BGCs (Figure 2b). Further analysis of the genomic context revealed that multiple modifying enzymes were well conserved in this family. We named this family the *mat* family and hypothesized that the additional oxidoreductase might introduce alternative modifications to the precursor peptide. The largest cluster in the SSN, which we termed the *sis* BGC family, also drew our interest. Similar to the *mat* family, the sequence logos of the *sis* family exhibited a characteristic C-terminal Avi(Me)Cys ring formation motif (T-X1-X2-X3-X4-C, T is threonine, C is cysteine, X represents a variable residue), and an additional conserved cysteine at the N-terminus (Figure 2c). However, analysis of the precursor sequence logo for the *sis* family revealed an approximately 50% occurrence of Cys residues in the middle of the core region, which could be further categorized into two subfamilies with either two or three Cys residues (Figure 2c). Considering the presence of these two precursor subfamilies within one BGC, we postulate that a single maturation system within the *sis* family could generate ACPs with diverse thioether rings, owing to the propensity of Cys residues to react with Dha or Dhb. Overall, our analysis prioritized potential BGC candidates for exploring the chemical diversity of Avi(Me)Cys-containing RiPPs.

**Figure 2.**
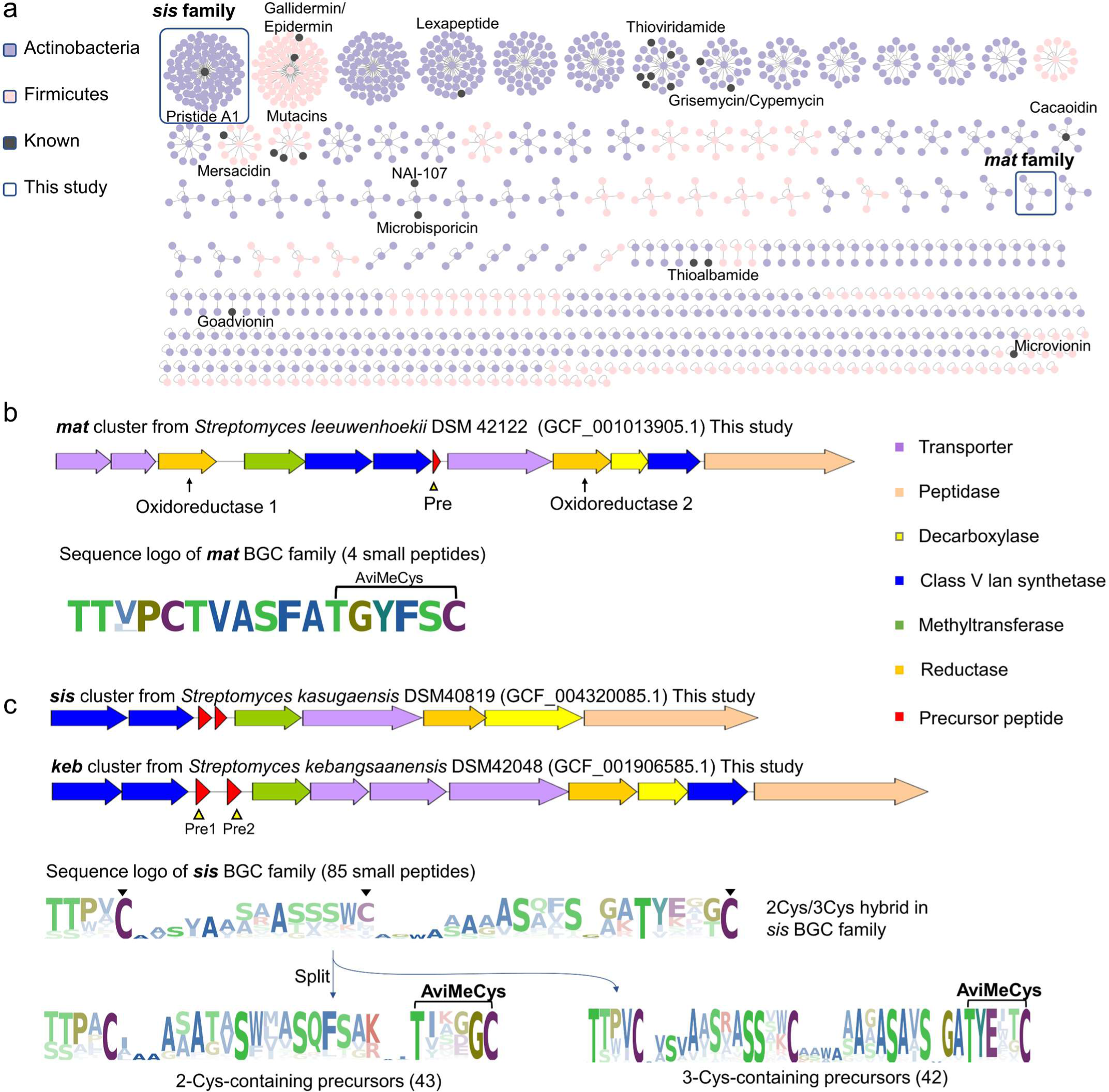
SPECO-based genome mining of ACP BGCs. a) A total of 1,172 precursor peptides were identified by SSN analysis, which are categorized to 67 clusters. The sequence logo of each cluster was shown in Supplementary dataset. Previously characterized precursors are highlighted as black dots. Two BGC families characterized in this study are boxed. b) BGC architecture and core peptide sequence logo of the *mat* BGC family. Oxidoreductase and precursor were highlighted by arrow and triangle, respectively. The black connector denotes Avi(Me)Cys ring formation. c) BGC architecture and core peptide sequence logo of the *sis* BGC family. Cys residues were highlighted by solid triangles.

### Linking Avi(Me)Cys biosynthetic gene clusters with peptide antibiotics

To discover new ACPs from BGCs of interest, we first focused on the *mat* cluster from *Streptomyces leeuwenhoekii* DSM42122 (Figure 3a). The *mat* cluster contains several enzymes responsible for the formation of the (methyl)lanthionine ((Me)Lan) ring, including MatKYX enzymes that share similarities with the class V lanthionine (Lan) synthetase LxmKYX from the lexapeptide BGC^26, 34^. Other enzymes in the cluster include MatM, which functions as a methyltransferase, and MatT1-T3 and MatP, which act as transporters and peptidase, respectively. Additionally, MatR1 and MatR2 are putative oxidoreductases involved in the hydrogenation of Dha or Dhb^26, 29^. The intricate modifications, such as decarboxylation, dehydration, thioether cyclization, hydrogenation, and methylation, give rise to a diverse range of possible monoisotopic masses for both intact and fragmented precursors (Figure 3b, Supplementary dataset). Consequently, targeting specific metabolites from the metabolic analysis of crude extracts can present challenges, as numerous *m/z* signals are observed, making it difficult to pinpoint the metabolites of interest.

**Figure 3.**
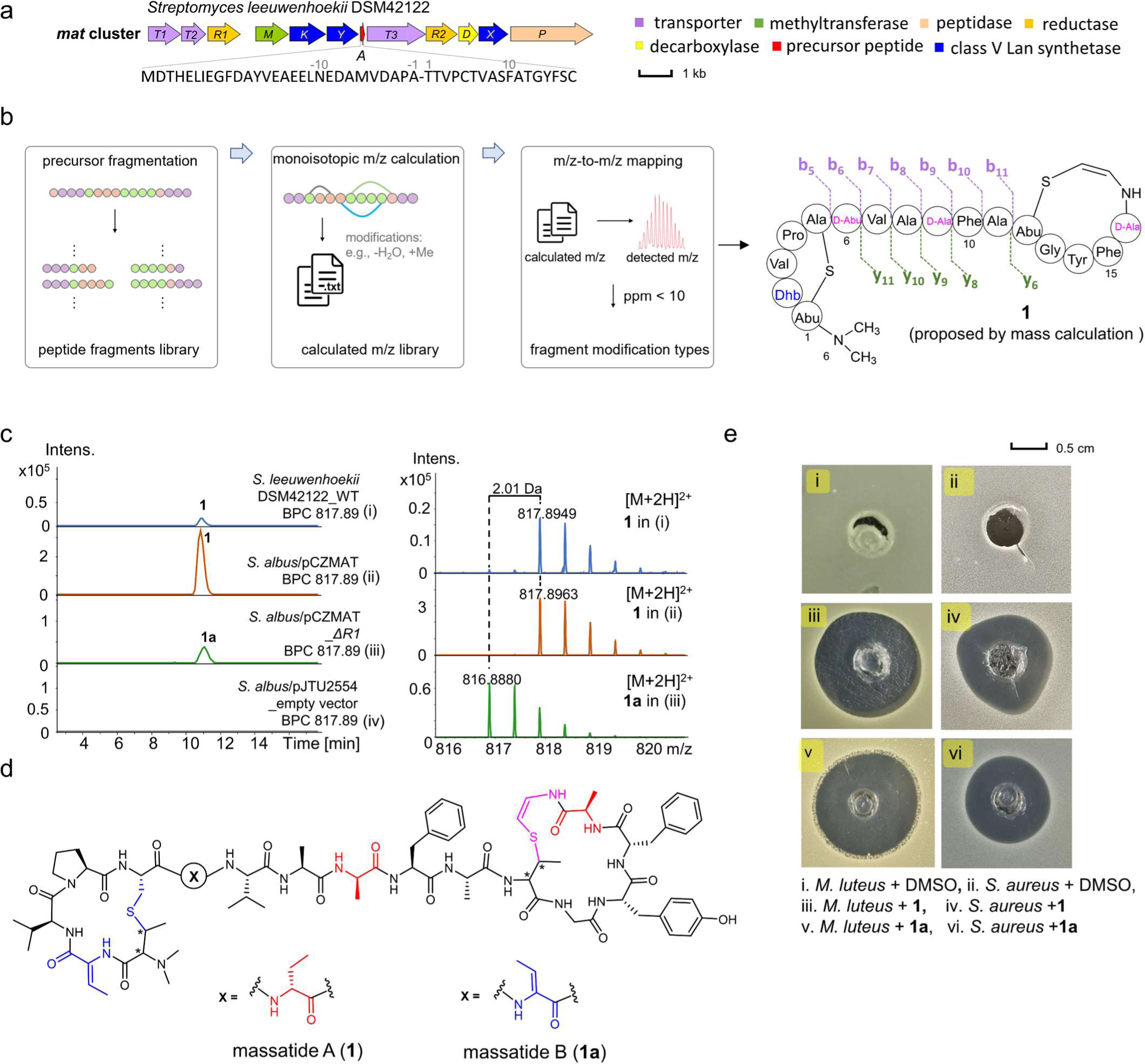
Discovery and BGC characterization of massatide. a) The *mat* BGC and amino acid sequence of the precursor MatA. b) Automatic mass mapping pipeline and proposed structure of **1** based on tandem mass analysis. Calculation details are in the Supplementary note. c) UPLC-HRMS analysis of *Streptomyces leeuwenhoekii* DSM42122 wild-type strain (i), *S. albus*/pCZMAT (ii, heterologous expression of *mat* BGC), *S. albus*/pCZMAT_*ΔR1* (iii, *matR1* deletion) and *S. albus*/pJTU2554 (iv, empty vector). For raw data, see Figure S2. d) The structure of massatide. AviCys motifs are shown in pink. D-amino acids are shown in red. Other noncanonical amino acids are shown in blue. e) Bioactivity of massatides against *S. aureus* and *M. luteus*.

To find the BGC-encoded ACP from metabolites, we matched calculated mass data with experimentally detected metabolomic data to target natural products (Figure 3b). This process involved enumerating *m/z* values of theoretical peptide fragments with different modifications and charge states and then matching them with experimental HRMS data. For instance, UPLC-HRMS analysis of the culture extract showed typical peptidic signals and a mass matching pipeline revealed a hit of [*M*+2H]^2+^=817.8949 (Figure 3b, Supplementary note, Supplementary dataset). Further mass calculation suggested that the core region of the precursor peptide “TTVPCTVASFATGYFSC” (calculated [*M*+2H]^2+^ = 817.8965, Δppm = 2.0) underwent six dehydrations (−6H_2_O), one decarboxylation (-CO_2_H_2_), three hydrogenations (+6H), and two methylations (+2CH_2_). To further link the metabolite of [*M*+2H]^2+^=817.8949 (**1**) with the *mat* BGC, we cloned and heterologously expressed the *mat* BGC in *Streptomyces albus* J1074. UPLC-HRMS analysis revealed a new peak with [*M*+2H]^2+^=817.8963 in the *S. albus*/pCZMAT by comparison with host strain harboring the empty pJTU2554 vector (Figure 3c, S2-S3). This result conclusively established that the *mat* BGC encodes compound **1**, with MatA as the crucial precursor for structural elucidation.

With the confirmed core peptide sequence and calculated modifications, we aligned experimental tandem mass data with calculated monoisotopic mass data of fragment ions (Figure S3). The results showed that experimental b5 to b11, y6, and y8 to y11 fragments with modifications matched well with calculated ones. The matched modifications at ions b11(−4H_2_O_+2CH_2__+4H) and y6 (−2H_2_O_-1CO_2_H_2__+2H) are consistent with the total of six dehydrations (−6H_2_O) and C-terminal decarboxylation. Comparing b5 to b6 and b8 to b9 ions localized two hydrogenated residues at Thr6 and Ser9, respectively. The absence of fragment ions in the Thr1-Cys5 and Thr12-Cys17 motifs strongly suggests the formations of Thr1-to-Cys5 linkage and Thr12-to-Cys17 linkage, respectively, with Ser16 being the only remaining residue that can be hydrogenated. It is noteworthy that hydrogenation during RiPP biosynthesis can potentially generate D-amino acids, as previously reported^26, 35, 36^. Based on the involvement of biosynthetic enzymes like methyltransferase and decarboxylase, we proposed a tentative structure for **1**, which we named massatide A (Figure 3b).

To further confirm the structure of compound **1**, a large-scale fermentation of the *S. albus*/pCZMAT strain was performed, and 5 mg of **1** was purified for 1D and 2D nuclear magnetic resonance (NMR) analysis (Figure S4-11, Table S4). In the ^1^H and ^13^C spectra, characteristic signals of the Phe and Tyr side chains and the modified residues Abu, Dhb, and AviMeCys were identified. A *Z*-geometry for the double bond of the AviMeCys residue was determined based on the corresponding ^3^*J*_H,H_ value of 8.5 Hz (Table S4). A C-S bond formation between AviMeCys and Thr12 was also supported by key HMBC correlations from AviMeCys-H to Thr12-Cβ and from Thr12-Hβ to AviMeCys-C. A Key HMBC correlation from Thr1-Hβ to Cys5-Cβ indicated a C-S crosslink between Cys-S and Thr-Cβ, consistent with MS/MS data. Furthermore, an *N,N*-dimethyl MeLan was observed based on HSQC and HMBC spectra, which is rarely reported except for a lanthidin RiPP^29^. To confirm absolute configurations for all the amino acid residues, we conducted advanced Marfey’s analysis^37^ for **1** using L/D-FDLA (5-fluoro-2,4-dinitrophenyl)-L/D-leucinamide). The results showed the presence of both L- and D-Ala, with D-Abu also observed (Table S8). Further investigation showed that Ala8 and Ala11 were genetically encoded and in the L-configuration. Based on these results, we proposed that Thr6, Ser9, and Ser16 were converted to D-Abu and D-Ala in **1**, while the remaining unmodified residues all existed as L-configuration. To the best of our knowledge, massatide A (**1**) represents the first Avi(Me)Cys-containing peptide with D-amino acids in the C-terminal AviMeCys ring.

Most reported lanthipeptide BGCs have only one reductase for Dha/Dhb hydrogenation^26, 35, 36^. However, in the *mat* BGC, both *matR1* and *matR2* encode LLM class oxidoreductases containing the conserved F420 binding domain, which belong to the LanJ_C_ family of lanthipeptide reductases^26^. To determine the essentiality of both MatR1 and MatR2, we constructed a knockout strain *S. albus*/pCZMAT_*ΔR1*. UPLC-HRMS data showed that compound **1a** ([*M*+2H]^2+^=816.8880) from the knockout strain, named massatide B, was 2 Da lighter than **1** (Figure 3c). Tandem mass analysis suggested that Thr6 was converted to Dhb in massatide B (**1a**) instead of D-Abu in **1** (Figures 1d, S12). Thus, we confirmed that in the *mat* BGC, MatR1 catalyzes D-Abu6 formation, while MatR2 may be responsible for both D-Ala9 and D-Ala16 formation.

### Two crosslinking patterns catalyzed by one Avi(Me)Cys biosynthetic machinery

We next investigated the *sis* family, which is predicted to catalyze two distinct crosslinking patterns, aiming to convert its biosynthetic potential into peptide antibiotics. We examined a *sis* BGC from *Streptomyces kasugaensis* DSM40819, which belongs to class V lanthipeptides with methylation and AviMeCys motif (Figure 4a, Table S3). The precursor genes *sis*A1 and *sis*A2, which encode small peptides containing two cysteines and three cysteines, respectively, could produce a two-ring and three-ring peptide. Using the same mass mapping workflow, we pinpointed two putative peptide ion peaks corresponding to the precursors SisA1(compound **2**, observed [*M*+3H]^3+^ = 1085.2083, calculated [*M*+3H]^3+^ = 1085.2083, Δppm = 0.1) and SisA2 (compound **3**, observed [*M*+3H]^3+^ = 816.7481, calculated [*M*+3H]^3+^ = 816.7487, Δppm = 0.7), respectively (Figure S13). Modifications of SisA1 occurred in the C-terminal 34-mer core region, including eleven dehydrations (−11H_2_O), one decarboxylation (-CO_2_H_2_), five hydrogenations (+10H), and two methylations (+2CH_2_), resulting in the production of compound **2**. Similarly, modifications of the SisA2 core peptide “STPACGAATVSWIVSQFSAKTVKDGC” involved seven dehydrations (−7H_2_O), one decarboxylation (-CO_2_H_2_), and three hydrogenations (+6H), leading to the production of **3** (Supplementary dataset). To validate the ribosomal origins of compounds **2** and **3**, we cloned and heterologously expressed the *sis* BGC in *S. albus.* Comparative UPLC-HRMS analysis revealed two new peaks in *S. albus*/pCZSIS, [*M*+3H]^3+^=1085.2076 and [*M*+3H]^3+^=816.7487, which are identical to compounds **2** and **3** from wild-type strain (Figures S13, S14, S23). The results confirmed that compounds **2** and **3**, named sistertide A1 and A2, were encoded by the *sis* BGC.

**Figure 4.**
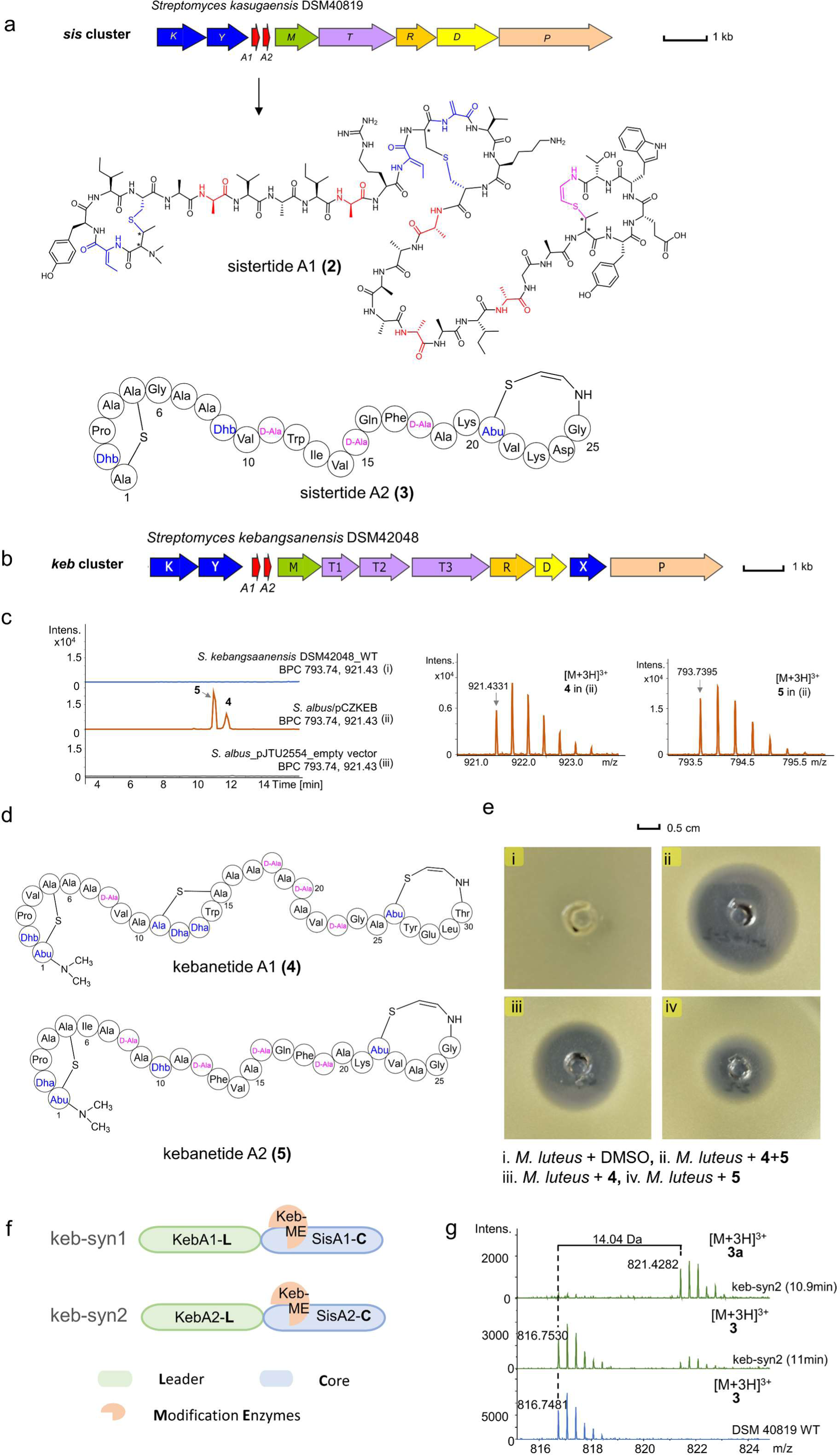
Discovery and characterization of the *sis* BGC family. a) The *sis* BGC and structures of sistertide A1 and sistertide A2. AviMeCys motifs are shown in pink. D-amino acids are shown in red. Other noncanonical amino acid are shown in blue. b) The *keb* BGC. c) UPLC-HRMS analysis of *Streptomyces kebangsaanensis* DSM42048 wild type strain (i), *S. albus*/pCZKEB (ii, heterologous expression of *keb* BGC) and *S. albus*/pJTU2554 (iv, empty vector). For raw data, see figure S24. d) Structure of kebanetides. e) Bioactivity of kebanetides against *M. luteus*. f) Chimeric precursor design. g) UPLC-HRMS analysis of constructs keb-syn2.

Tandem mass analysis of sistertide A1 (**2**) fragments revealed N-terminal methylation and C-terminal decarboxylation (Figure 4a, S14). Furthermore, the analysis indicated that Ser7, Ser11, Ser19, Ser23, and Ser26 underwent a 16 Da mass loss, suggesting that they were first dehydrated (-H_2_O) and then reduced (+2H). Notably, no fragment ions were observed around three cysteines, which implied that Cys5, Cys18, and Cys34 might cyclize with Thr1, Ser14, and Thr29 (Figures 4a, S14). We next performed a large-scale fermentation of *S. albus*/pCZSIS to determine the crosslinking patterns and obtained 4 mg pure sistertide A1 (**2**) for NMR analysis (Figures S15-22, Table S5). Characteristic signals of the side chains of Tyr, Trp, Thr, and modified residues Dha and Dhb were observed in **2**. NMR analysis also showed that **2** contains an AviMeCys scaffold with a *Z*-geometry (^3^*J*_H,H_ 8.4 Hz) at the double bond, which was derived from the Thr29/Cys34 motif of the core peptide (Table S5). The presence of *N,N*-dimethyl MeLan was confirmed by key HMBC correlations (Figures S19, 22). However, due to its low yield, we could not collect enough sistertide A2 (**3**) for NMR analysis. Mass calculation and tandem mass data of **3** suggested that the core peptide of SisA2 underwent dehydration, decarboxylation, hydrogenation, and ring formations, which led to the formation of one lanthionine ring and one AviMeCys ring (Figures 4a, S23). Sistertide A1 is methylated, while A2 lacks this modification despite being modified by the same enzymes. This suggests that the methyltransferase in the *sis* BGC may exhibit selectivity in substrate recognition. The presence of a putative oxidoreductase SisR in this BGC suggests possible D-amino acid formation in sistertides. Advanced Marfey’s analysis^37^ for compound **2** revealed the presence of both L- and D-Ala in the structure (Table S9), with the D-Ala residues likely converted from Ser7, 11, 19, 23, and 26 in the precursor. The remaining unmodified residues were L-configurated. Similarly, for sistertide A2 (**3**), Ala11, Ala15, and Ala18 were likely resulted from dehydration and reduction of Ser residues in the precursor, suggesting that they may also be D-Ala in the mature compound based on biosynthetic logic.

Additionally, within the *sis* family, we identified two other ACPs encoded by a *sis* BGC homolog, the *keb* cluster from *Streptomyces kebangsaanensis* DSM42048 (Figure 4b). Our initial attempt to identify target ACPs from the fermentation extract of the wild-type DSM42048 was unsuccessful. Subsequently, we tried to heterologously express the entire *keb* BGC in *Streptomyces albus*, which resulted in the production of two new compounds, **4** (named kebanetide A1, [*M*+3H]^3+^ = *m/z* 921.4331) and **5** (named kebanetide A2, [*M*+3H]^3+^ = *m/z* 793.7395) (Figures 4c, S24). Using mass calculation, tandem mass spectrum analysis, ^1^H NMR spectroscopy, and advanced Marfey’s analysis, we determined that kebanetide A1 (**4**) features one C-terminal AviMeCys ring and two thioether crosslinks. Kebanetide A2 (**5**) resembles sistertide A2, containing a C-terminal AviMeCys and N-terminal thioether ring (Figures 4d, S25-32, Table S6-S7, S10-S11).

Although the leader peptide remains conserved across multiple precursors within the *sis* family, significant variations are observed in the core regions of precursor A1 and A2, particularly in terms of their length and the number of essential cysteine residues involved in ring formation. Notably, the AviMeCys ring of sistertide A1 is characterized by the presence of two bulky hydrophobic aromatic side chains (Try30 and Trp32) and one negatively charged residue (Glu31). On the other hand, sistertide A2 contains a positively charged lysine residue (Lys23). These significant differences in the core peptide suggest that the tailoring enzymes within the *sis* BGC family exhibited broad substrate selectivity. It is worth noting that the discovery of 42 BGCs in this family indicates that the “one tailoring enzyme system and multiple crosslinking patterns” are widespread among actinobacteria. RiPP biosynthetic cassettes with multiple precursors were found in two-component class II lantibiotics, such as lichenicidin^38^ and haloduracin^39^. However, unlike the *sis* family, multiple LanM enzymes were encoded in these two-component lantibiotics BGCs to catalyze precursors in a one-LanM-one-precursor manner. Ribosomal peptide maturases in the *sis* family resemble promiscuous lanthipeptide synthetase ProcM, which has been widely used to generate cyclic peptide libraries^40^.

To explore the catalytic potential of the *sis* family maturase, we conducted a leader-core mixing and matching assay. We designed two chimeric precursor peptides that combined the KebA leader with the SisA core region, respectively, to investigate the substrate tolerance in ACP biosynthesis (Figures 4f, S33). Co-expression of these two chimeric precursors with the maturases from the *keb* BGC (constructs keb-syn1-2) led to the production of compound **2a,** sistertide A2 (**3**) and **3a** (Figures 4g, S34-35). Tandem mass analysis supported that **3a** is monomethylated **3** at the N-terminus (Figure S35), indicating the *keb* maturase can not only modify the core peptide of SisA2, but also diversify the end product via methylation. HRMS data showed that **2a** is a new derivative compared to sistertide A1 (**2**). Further tandem mass spectrometry analysis indicated that the mass differences were localized in the N-terminus (Figure S34). No new peaks were observed when chimeric precursors Syn1 and Syn2 coexpressed with the *sis* maturases (constructs sis-syn1-2), implying leader-maturase mismatching hindered core peptide modification. Taken together, these findings suggested that the tailoring enzymes from the *keb* BGC exhibited leader peptide selectivity but core peptide tolerance.

### Potent antibacterial activity of ACPs

We next employed the Kirby-Bauer assay to evaluate the antibacterial properties of isolated ACPs of the *mat* and *sis* families. Massatides A (**1**) and B (**1a**) showed strong inhibitory activity against *Staphylococcus aureus* and *Micrococcus luteus* (Figure 3e). Similarly, kebanetides A1 (**4**) and A2 (**5**) inhibited the growth of *M. luteus* as shown in Figure 4e. Furthermore, we investigated the potential synergy between kebanetide A1 and A2, as two-component lantibiotics typically function by binding to lipid II and the compound-lipid II complex^41^. The results indicated a slightly larger inhibition zone in Figure 4e, suggesting a weak synergistic effect between kebanetide A1 and A2.

Moreover, we determined the minimal inhibitory concentrations (MIC) of massatide A, massatide B, sistertide A1, kebanetide A1, and A2 against seven bacterial strains to assess their bioactivity, as detailed in Table 1. Massatide A exhibited a broad spectrum of efficacy against gram-positive bacteria, with lower MICs than nisin and comparable activity to vancomycin. Its antimicrobial activities against *S. aureus* and *M. luteus* even surpassed those of vancomycin, highlighting its potential as a promising candidate for gram-positive antibiotics. The activity of massatide B was similar to that of massatide A, with only a slightly larger MIC, indicating that the presence of D-Abu6 has a minimal contribution to the antibacterial activity. On the other hand, sistertide A1 alone did not exhibit inhibitory activity against the tested strains. Kebanetide A1 and A2 demonstrated weaker bioactivity than massatides but performed better than nisin against *E. faecalis* and *E. faecium*. A possible synergistic effect was observed when kebanetide A1 and A2 were combined at a 1:1 ratio. No bioactivity was observed against gram-negative bacteria. The growth kinetics and time-dependent killing assay revealed that massatide A was bactericidal against *S. aureus* and was superior to vancomycin, the first line of defense antibiotic, in killing early exponential phase populations (Figure 5a,c). We noticed that massatide A did not result in lysis of the cell culture, which is the same as vancomycin (Figure 5b).

**Table 1.** MIC values of massatides, sistertide A1 and kebanetides (μg/mL).

| strains |  | massatide<br>A (1) | massatide<br>B (1a) | sistertide<br>A1 (2) | kebanetide<br>A1 (4) | kebanetide<br>A2 (5) | kebanetides<br>(4+5) | nisin | vancomycin |
| --- | --- | --- | --- | --- | --- | --- | --- | --- | --- |
| G+ | *Staphylococcus aureus* ATCC25923 | 0.5 | 1 | >64 | 64 | >64 | 64 | 64 | 1 |
|  | *Micrococcus luteus* DSM1790 | 0.06 | 0.25 | >64 | 8 | 32 | 8-16 | 0.5 | 0.125 |
|  | *Bacillus subtilis* 168 | 2 | 4 | >64 | 8 | 32-64 | 16 | 32 | 0.125 |
|  | *Enterococcus faecalis* OG1RF | 4 | 4 | >64 | 16 | 16 | 8 | >64 | 2 |
|  | *Enterococcus faecium* MCC2763 | 2 | 4 | >64 | 32 | 32 | 16 | >64 | 1 |
| G- | *Escherichia coli* DH5α | >128 | >64 | >64 | >64 | / | / | / | / |
|  | *Pseudomonas aeruginosa* PAO1 | >128 | >64 | >64 | >64 | / | / | / | / |

**Figure 5.**
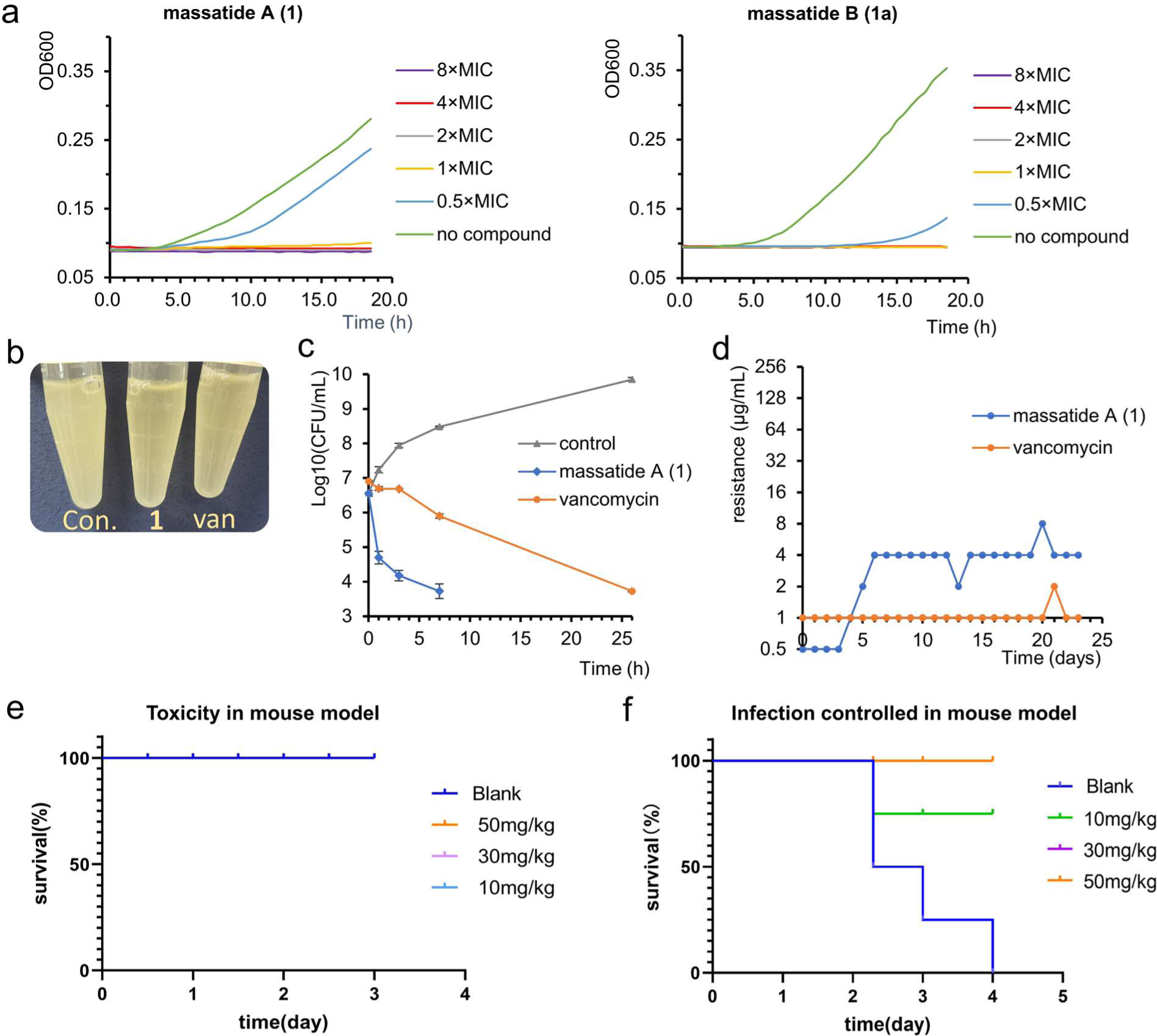
Bioactivity of massatide. a) The growth kinetics of *S. aureus* with massatide A and massatide B. b) Massatide A did not resulted in lysis. c) Time-dependent killing of *S. aureus* (10×MIC vancomycin, 10×MIC massatide A, 10×MIC massatide B). d) Resistance acquisition during serial passaging in the presence of sub-MIC levels. e)The survival rate in acute toxicity study *in vivo*. f) The survival rate in septicemia model using MRSA treated with massatide A.

In light of the promising antibacterial potential of massatide A, we conducted extensive research to investigate its antibacterial efficacy both *in vitro* and *in vivo*. The subsequent antibacterial assays yielded remarkable results, demonstrating that massatide A exhibited excellent activity against a range of clinically isolated resistant pathogens, as outlined in Table 2. Its potency against tested gram-positive pathogens, including vancomycin intermediate-resistant strains, remained below 4 μg/mL. An outstanding finding was the exceptional activity of massatide A against linezolid-resistant *S. aureus* 20-1 and methicillin-resistant *S. aureus* ATCC43300, with a minimum inhibitory concentration (MIC) of only 0.25 μg/mL. During the process of serial passaging in subinhibitory concentrations of massatide A, there was only a slight increase in the minimum inhibitory concentration (MIC) to *S. aureus*, from 0.5 to 4 μg/mL, over a span of 23 days. No significant change in MIC was observed during the subsequent 17 days after day 6, suggesting that *S. aureus* did not develop resistance to this compound. This finding further supports the potential of massatide A as an effective antibacterial agent.

**Table 2.** Biological activity of massatide A against clinic isolated resistant strains (μg/mL).

| strains |  | massatide A(1) | levofloxacin |
| --- | --- | --- | --- |
| G+ | MRSA 15-1 | 2 | 16 |
|  | LRSA 20-1 | 0.25 | 32 |
|  | MRSA ATCC33591 | 1 | 0.25 |
|  | MRSA ATCC43300 | 0.25 | 0.25 |
|  | VMSA ATCC700788 | 4 | 32 |
|  | VMSA ATCC700699 | 4 | 64 |
|  | *Enterococcus faecalis* 19-1 | 4 | 1 |
|  | LREfa 20-1 | 4 | 32 |
|  | VREfa ATCC51575 | 4 | 0.5 |
| G- | *Acinetobacter baumannii* 20-1 | 64 | 8 |
|  | *Acinetobacter baumannii* 20-2 | 64 | 0.06 |
|  | *Klebsiella pneumoniae* ATCC700603 | >64 | 0.5 |
|  | *Klebsiella pneumoniae* 15-1 | >64 | 0.25 |

\*MRSA: Methicillin-resistant *Staphylococcus aureus*; LRSA: Linezolid-resistant *Staphylococcus aureus*; VMSA: vancomycin-intermediate *Staphylococcus aureus*; LREfa: Linezolid-resistant *Enterococcus faecalis*; VREfa: Vancomycin-resistant *Enterococcus faecalis*.

Furthermore, massatides exhibited no cytotoxicity against Hela cells at a 200 μg/mL concentration but demonstrated moderate toxicity against the human cell line Hek293T (Figure S36). To further assess its safety profile, an acute toxicity study was conducted in mice, where massatide A was administered at single doses ranging from 10-50 mg/kg. After 72 hours of injection, all treated mice survived without any signs of acute toxicity (Figure 5e). These findings suggest massatide A has a favorable safety profile and highlight its potential as a safe and effective antibacterial agent.

Expanding upon its potent antibacterial activity, low *in vivo* toxicity, and minimal risk of resistance, we conducted further evaluations to assess the efficacy of massatide A in a mouse septicemia model. Mice were intraperitoneally infected with 7.2 × 10^9^ c.f.u. of MRSA ATCC43300, and lethality was observed after 96 hours of infection (Figure 5f). Remarkably, the survival rate of mice significantly improved to 80% at a single dose of 10 mg/kg of massatide A, and further increased to 100% at a higher dosage of 50 mg/kg, when compared to the control group with no survival. These findings strongly indicate the effectiveness of massatide A *in vivo* as a potential therapeutic agent. Taken together, these results highlight the immense potential of massatide A for development as a therapeutic agent against gram-positive pathogens. Its remarkable efficacy and favorable safety profile make it a promising candidate for further research and development in the field of antibacterial treatment.

## Discussion

Bacterial ribosomal peptides have long been recognized as a valuable source for antibiotic discovery^42^. However, identifying antibiotic peptides remains challenging due to high rediscovery rates. Recent genome sequencing of bacteria has revealed an untapped reservoir of biosynthetic gene clusters (BGCs), but the challenge remains in prioritizing BGCs of interest and robustly linking metabolites to their corresponding BGCs before time-consuming isolation. In this study, we addressed these challenges by leveraging the genetically-encoded nature of RiPPs to expand the diversity of Avi(Me)Cys-RiPPs for antibiotic discovery. Our comprehensive approach involved integrating rule-based genome mining, automated metabolomic analysis, and heterologous expression, enabling us to link new BGCs with their encoding ACP chemicals directly. Through this approach, we successfully discovered five ACPs that exhibited potent antibacterial activity against a wide range of gram-positive pathogens. However, our approach may have missed novel compounds with unknown modifications that were not predictable by automated metabolomic analysis. Among the uncharacterized BGCs in our dataset, we identified a few containing a DUF3105 domain-containing protein (Figure S37), suggesting the presence of an unknown modification that could be installed on the precursor. Improving metabolomics analysis strategies, including mass calculation and mapping, could help alleviate such situations. Nonetheless, our study demonstrates the potential of a comprehensive approach in identifying novel antibiotic peptides from the underexplored RiPP families.

Flavoprotein, the essential enzyme in ACP BGCs, is widespread in bacterial genomes (Figure S38). By using the SPECO genome mining pipeline, we identified 1,172 BGCs from actinobacteria and firmicutes genomes, representing 67 ACP families for potential antibiotic discovery. These diverse ACP BGCs, in combination with the widely distributed flavoprotein in bacteria, represent an underexplored source for exploring the chemical diversity of Avi(Me)Cys-containing RiPPs, indicating their potential as a promising untapped source for genomics-guieded discovery of peptide antibiotic. For instance, in addition to the known ACP families, a recent finding by Cheng et al. was reported that flavoproteins could couple with radical S-adenosylmethionine enzymes to catalyze a new unsaturated thioether residue, S-[2-aminovinyl]-3-carbamoylcysteine (AviCamCys)^43^. Whether the compound with this new structure showed biological activity remained unknown. The discovery of flavoproteins in different families of RiPPs highlights the potential of exploring the untapped sequence space of flavoproteins for discovering novel RiPPs.

Our genomics-guided discovery of underexplored ACPs led to the identification of massatide A, a peptide antibiotic that demonstrated potent antibacterial activity both *in vitro* and *in vivo*, with low toxicity and minimal risk of resistance. These findings highlight the significant potential of ACPs as therapeutic candidates against gram-positive pathogens, making them a promising candidate for further development in antibacterial treatment. However, further structural modifications are necessary to enhance its drug-like properties, including improvements in water solubility and safety profile, particularly concerning human kidney cells. The *sis* family of ACPs serves as an excellent example of the substrate tolerance of ACP biosynthesis, providing a promising avenue for the bioengineering of new-to-nature analogues with desired properties and the development of novel antibiotics. Despite the need for further optimization to improve the yield and scalability of the heterologous expression system, the substrate tolerance of ACP biosynthesis remains a promising approach for peptide bioengineering and the advancement of new antibiotics. We believe that leveraging the promiscuity of ACP biosynthesis will enable the creation of effective antibiotics with desired properties, including simple structures, high efficacy, and low toxicity. These modifications will pave the way for the advancement of ACP antibiotics as a safer and more effective antibacterial agent in the future.

Our workflow, guided by omics analysis, provides a promising strategy for the targeted discovery of RiPPs with antibiotic activity. We envision that our efforts in genome mining, characterization, and bioactivity study of ACPs will contribute to the discovery of new RiPP natural products, expanding the repertoire of compounds available for antibiotic development. This could potentially address the urgent need for novel antibiotics to combat the emergence of antibiotic-resistant bacteria.

## Acknowledgments

This work is funded by the Research Grants Council of Hong Kong (27107320, 17115322, and 17102123) to Y.X. Li and the CAMS Innovation Fund for Medical Sciences (2021-I2M-1-030) to W. Z.. The authors would like to thank Prof. Meifeng Tao for the gift of pJTU2554 vector and Dr. Prasanna Neelakantan for the gift of indicator strains. We also thank Yi-Man Eva Fung, Jo Yip and Bonnie Yan for their help in MS and NMR analysis.

## Author contributions

Y.X. L., X.Y. T., W. Z, Z. C. and B.B. H. designed the project and prepared the manuscript. Z. C. performed experiments and MS data analysis. B.B. H. performed bioinformatic analysis. K. L. conducted the animal experiments under the supervision of W. Z., and S. W.. Y. G. contributed to the mass mapping pipeline. H.Y. L. and R.Z. L. contribute to the NMR data analysis. Y.X. L. supervised the project.

## Conflicts of interest

The authors declare no conflicts of interest.

## Additional information

Supplementary information

Supplementary dataset

## Notes

### Competing Interest Statement

The authors have declared no competing interest.

